# Restoration of striatal neuroprotective pathways by kinase inhibitor treatment of Parkinson’s linked-LRRK2 mutant mice

**DOI:** 10.1101/2024.07.31.606089

**Authors:** Ebsy Jaimon, Yu-En Lin, Francesca Tonelli, Odetta Antico, Dario R. Alessi, Suzanne R. Pfeffer

## Abstract

Parkinson’s disease-associated, activating mutations in Leucine Rich Repeat Kinase 2 (LRRK2) block primary cilia formation in cholinergic and parvalbumin interneurons and astrocytes in the striatum, decreasing the production of GDNF and NRTN neuroprotective factors that normally support dopaminergic neuron viability. We show here that 3 month-dietary administration of the MLi-2 LRRK2 kinase inhibitor restores primary cilia and the Hedgehog-responsive production of neuroprotective GDNF and NRTN by these neurons; cilia are also restored on cholinergic neurons of the pedunculopontine nucleus. Importantly, we detect recovery of striatal dopaminergic processes and decreased stress-triggered Hedgehog signaling by nigral dopaminergic neurons. Thus, pathogenic LRRK2-driven cilia loss is reversible in post-mitotic neurons and astrocytes, which suggests that early administration of specific LRRK2 inhibitors may have significant therapeutic benefit for patients in the future.

**One Sentence Summary:** Kinase inhibitor restores cilia, Hedgehog signaling, neuroprotective factors and dopamine processes in Parkinson’s linked-LRRK2 mouse striatum

## INTRODUCTION

While the cause of most Parkinson’s disease (PD) is unknown, about 25% of cases are linked to genetic mutations, and activating mutations in the Leucine Rich Repeat Kinase 2 (LRRK2) are among the most common monogenic causes of PD (*1, 2*). LRRK2 phosphorylates a subset of Rab GTPases (*3, 4*) that are critical regulators of membrane trafficking (*5*). We have shown that LRRK2-mediated Rab phosphorylation blocks the formation of primary cilia in most cell types in culture, but only in specific cell types in mouse and human brain (*6–9*). In the striatum that is critical for dopamine signaling, LRRK2 mutation causes loss of cilia in a significant fraction of cholinergic and Parvalbumin interneurons and astrocytes, but no loss is seen in the much more abundant medium spiny neurons (*6–9*). Because cilia are essential for proper Hedgehog signaling (*10, 11*), cilia loss in the brain correlates with decreased Hedgehog signaling (*6, 8, 9*). We have shown that the consequence of cilia loss for cholinergic and Parvalbumin neurons is decreased production of the neuroprotective GDNF and NRTN peptides that support stressed dopamine neurons (*7, 9*). Thus, by blocking neuroprotective pathways, pathogenic LRRK2 will impact the vulnerability of dopamine neurons that are lost in PD.

Cilia are normally formed after cell division and disassembled in conjunction with the cell cycle (*12, 13*). The neurons and astrocytes that lose cilia in LRRK2 mutant brains are post-mitotic, non-dividing cells, thus it was not clear whether this process could be reversed by LRRK2 kinase inhibitors. Indeed, in our previous work, we did not see any rescue of cilia loss phenotypes after feeding mice with the MLi-2 LRRK2 inhibitor for two weeks (*8*). This is a very important issue, as multiple LRRK2 inhibitor clinical trials are currently underway (*14*). We show here that longer (3 month) dietary administration of MLi-2 reverses all ciliogenesis, Hedgehog signaling and neuroprotective peptide production phenotypes and restores cilia on cholinergic neurons of the pedunculopontine nucleus.

## RESULTS

### Rescue of primary cilia in striatal cholinergic neurons and astrocytes

We showed previously that striatal cholinergic neurons of mice with activating LRRK2 pathway mutations are ∼40% ciliated compared with ∼70% ciliated for wild type mice, sampling mice as young as 10 weeks and as old as 13 months (*6, 8*). In postmortem striatum of ∼85 year old patients, these neurons were ∼4% ciliated in both patients with idiopathic and LRRK2 pathway mutations compared with ∼12% for controls (*7*). Thus, it was important to determine whether ciliary defects can be reversed upon LRRK2 inhibitor administration. For these experiments, one group of mice received a modified rodent diet containing MLi-2 formulated to provide a concentration of 60 mg/kg per day based on the amount of diet that mice were estimated to consume. Previous work by Merck had demonstrated that this dose of MLi-2 administered via the diet results in full target engagement in the brain as monitored by pS935-LRRK2 levels (*15*), consistent with full LRRK2 inhibition. Mice were administered the ± MLi-2 diet for 3 months, and brains harvested at the same time of day in Scotland were analyzed for ciliation status in California before genotype identities were revealed. Immunoblot analysis of brain and kidney tissue confirmed marked decreased levels of pS935-LRRK2, pS105-Rab12 and pT73-Rab10 in the brain as well as kidney of animals administered the MLi-2 diet (Fig. 1A).

**Fig. 1.**
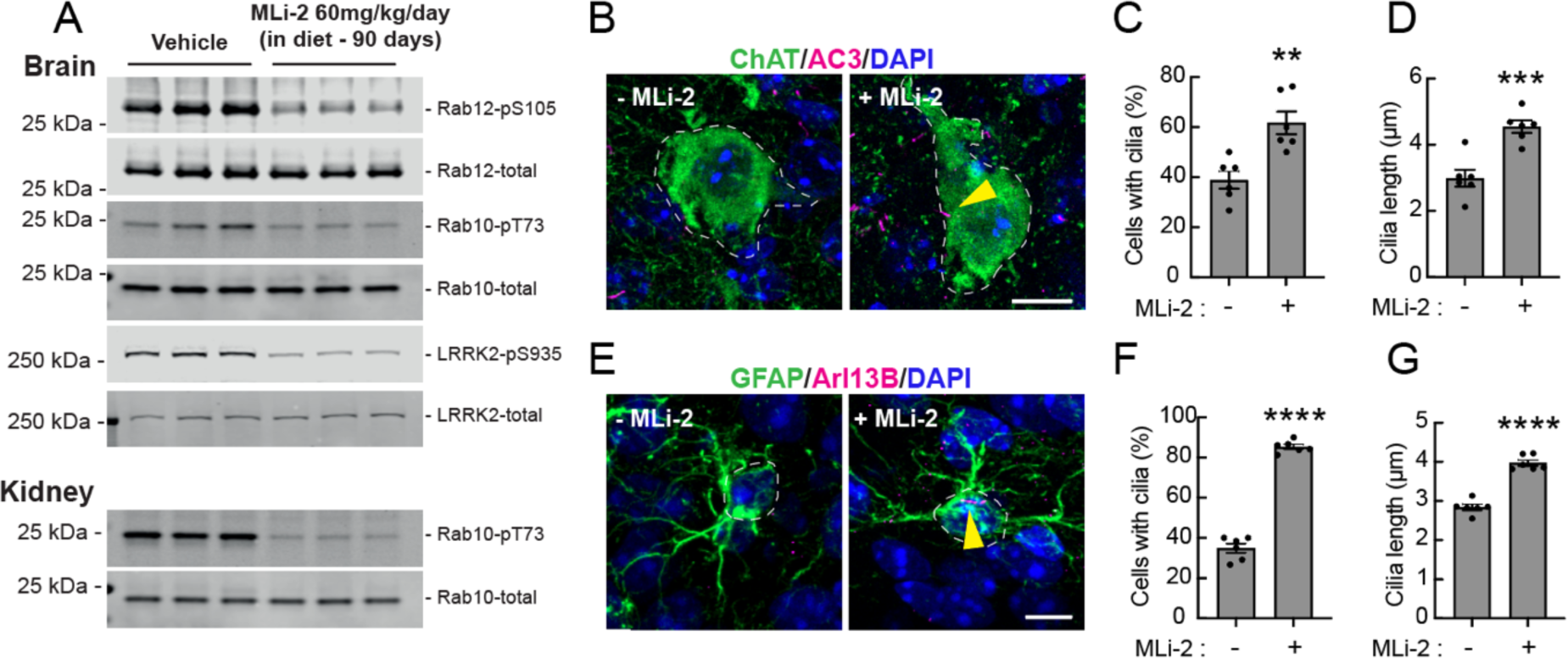
MLi-2 reverses cilia loss in mouse R1441C LRRK2 striatal cholinergic interneurons and astrocytes. (A) Quantitative immunoblots of mouse brain (upper) and kidney (lower) to monitor MLi-2 inhibition of LRRK2 after 90 days feeding, 60mg/kg/day. Each lane represents 12.5µg tissue from a different animal. (B) Confocal images of dorsal striatum from R1441C LRRK2 mice fed ± MLi-2 inhibitor for 3 months. Cholinergic interneurons were labeled using anti-ChAT antibody (green); primary cilia were labeled using anti-AC3 antibody (magenta; yellow arrowhead). Nuclei were labeled using DAPI (blue). (C) Quantitation of ChAT^+^ neurons containing a cilium and (D) cilia length. (E) Confocal images of astrocytes labeled using anti-GFAP antibody (green) and cilia labeled using anti-Arl13b (magenta, yellow arrowhead). (F) Quantitation of GFAP^+^ astrocytes containing a cilium and (G) cilia length. Error bars represent SEM from 6 R1441C LRRK2 brains ± MLi-2 with >30 cholinergic interneurons and astrocytes scored per brain. Statistical significance was determined using an unpaired t-test. Percentage ciliation in cholinergic interneurons: **p = 0.0027 for + MLi-2 versus - MLi-2 and cilia length in cholinergic interneurons: ***p = 0.0006 for + MLi-2 versus - MLi-2. Percentage ciliation in astrocytes: ****p<0.0001 for + MLi-2 versus - MLi-2 and cilia length in astrocytes: ****p<0.0001 for + MLi-2 versus - MLi-2. Bar, 10µm.

As shown in Figures 1B-D, in 5-month-old mice fed MLi-2 for 3 months, the percentage of ciliated cholinergic neurons was remarkably restored to almost wild type levels with two animals achieving wild type ciliation levels; cilia length also increased, which may confer increased signaling or inter-neuronal communication capacity to those cells (*16, 17*).

GFAP^+^ striatal astrocytes from LRRK2 G2019S knock-in mice (13-month-old, (*8*)) are ∼38% ciliated compared with ∼65% in wild type littermates. In 5-month-old LRRK2 R1441C mice, these astrocytes were also ∼38% ciliated but this reverted to >80% ciliated after 3 months of MLi-2 feeding (Fig. 1E,F). Moreover, the astrocyte cilia also increased in length (Fig. 1G). These experiments demonstrate that cilia can be restored to striatal cells upon MLi-2 administration.

### Cilia rescue in the pedunculopontine nucleus

Loss of cholinergic neurons in the pedunculopontine nucleus is linked to the severity of motor disability in PD patients (*18, 19*). Electrical stimulation of this region can produce organized locomotor movements, suggesting a direct role in movement initiation and control. We found that cholinergic neurons in the pedunculopontine nucleus are normally ∼70% ciliated; ciliation decreased significantly to ∼45% in 4-month-old LRRK2 R1441C animals (Fig. 2A,B). Cilia length also decreased from ∼4.5µm to ∼3µm (Fig. 2C). Due to cilia loss, these cells are expected to display a clear Hedgehog signaling deficit. After 3 months MLi-2 feeding, ciliation of these cholinergic neurons in 5-month-old mice improved from 40% to 70%, and length also increased (Fig. 2D-F). Thus, MLi-2 reverses ciliation defects in cholinergic neurons in the pedunculopontine nucleus as efficiently as in the striatum.

**Fig. 2.**
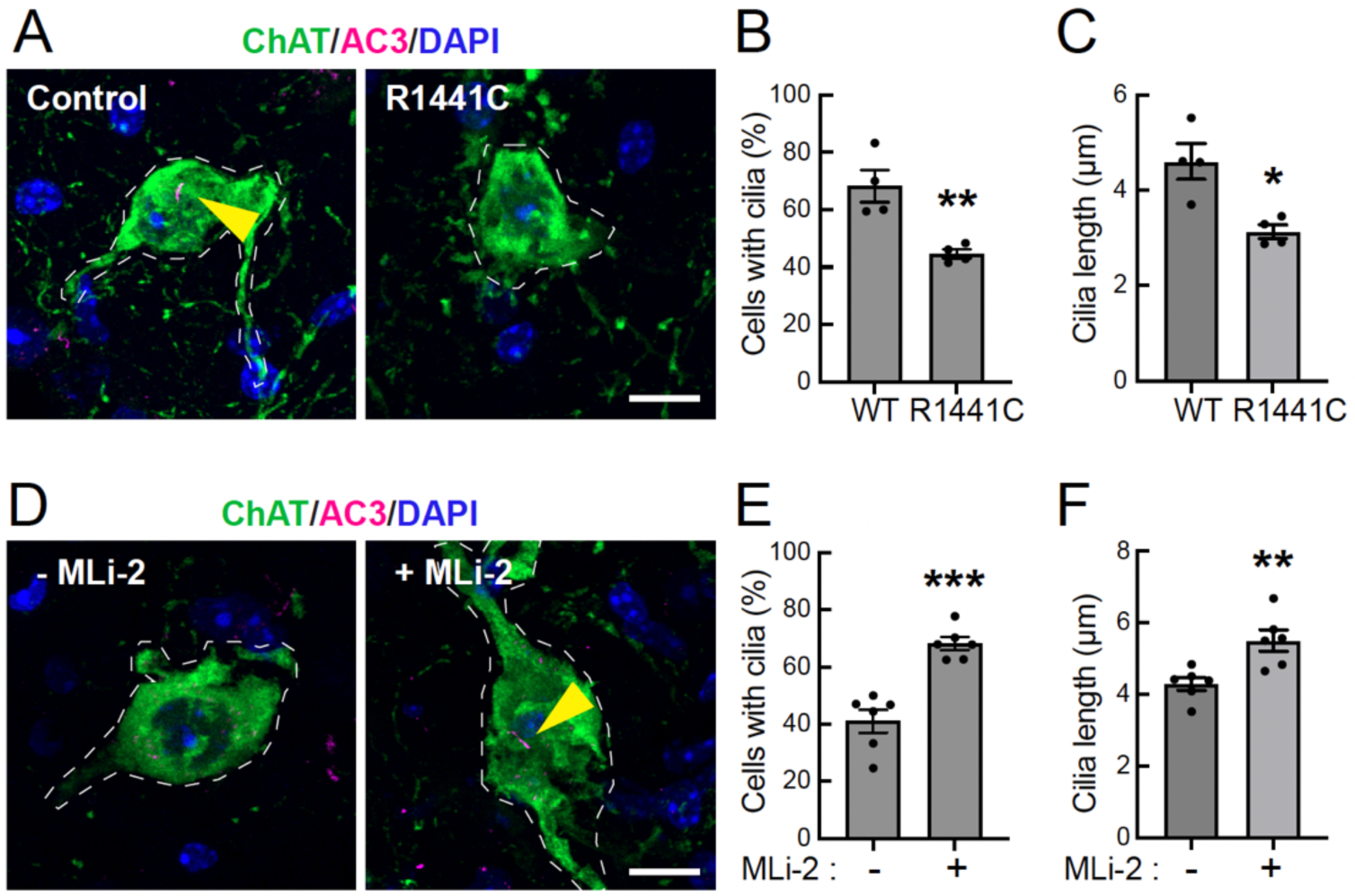
MLi-2 reverses cilia loss in mouse R1441C LRRK2 pedunculopontine nucleus (PPN) cholinergic neurons. (A) Confocal microscopy of the sections of the pedunculopontine nucleus (PPN) from 4-month-old WT and R1441C LRRK2 mice. Cholinergic neurons were labeled as in Fig. 1. (B) Quantitation of the percentage of ChAT^+^ neurons containing a cilium and (C) their cilia length. Error bars represent SEM from 4 R1441C LRRK2 brains with >30 cholinergic neurons scored per brain. (D) Similar analysis of PPN cholinergic neurons from R1441C LRRK2 mice fed with or without MLi-2 inhibitor-containing chow for 3 months. (E) Quantitation of the percentage of ChAT^+^ neurons containing a cilium and (F) their cilia length. Error bars represent SEM from 6 R1441C LRRK2 brains ± MLi-2 with >50 cholinergic neurons scored per brain. Statistical significance was determined using an unpaired t-test. Percentage ciliation in PPN cholinergic neurons: **p = 0.0068 for R1441C versus WT, Cilia length in PPN cholinergic neurons: *p = 0.0102 for R1441C versus WT, ***p = 0.0002 for + MLi-2 versus - MLi-2. **p = 0.0060 for + MLi-2 versus - MLi-2. Bar, 10µm.

### Restoration of Hedgehog signaling in striatal cholinergic interneurons and astrocytes

As mentioned earlier, cilia are needed for canonical Hedgehog signaling (*10–12*). We used RNAscope fluorescence in situ hybridization to monitor Hedgehog-dependent induction of Patched 1 (PTCH1) expression, an established marker of Hedgehog pathway signaling. Figure 3A shows example images of cholinergic neurons from 5-month-old wild type and LRRK2 G2019S mutant animals, with and without cilia. Quantitation of these images revealed that LRRK2 mutant animals showed ∼50% decrease in PTCH1 RNA levels (Fig. 3B). Control reactions carried out in the absence of an RNAscope probe did not show any RNA signal (Supplemental Fig. 1). In wild type cholinergic interneurons, ciliated cells showed the highest expression of PTCH1; this was also true in the mutant cells, except that even ciliated LRRK2 mutant cholinergic neurons expressed less PTCH1 than the level seen in wild type brains (Fig. 3C). Five-month-old LRRK2 R1441C mice expressed slightly less PTCH1 than the LRRK2 G2019S animals but this increased about 4-fold upon MLi-2 administration (Fig. 3D,E). Again, the highest expression was seen in ciliated cells upon drug treatment, and rescue reached the level seen in wild type control mice (Fig. 3F). Ciliated cholinergic neurons also showed decreased PTCH1 expression, indicating a signaling defect in both G2019S and R1441C LRRK2 mouse striatum.

**Fig. 3.**
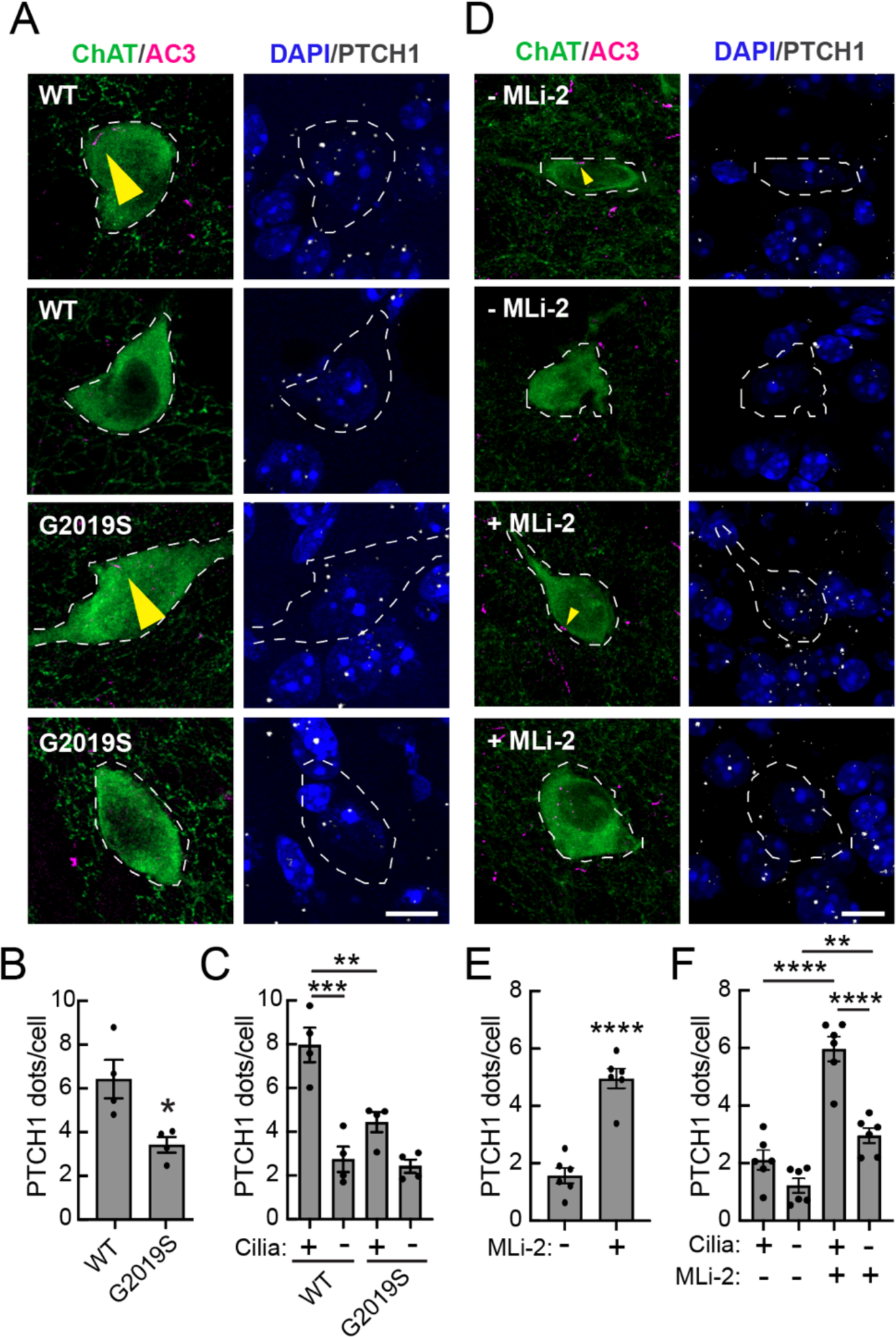
MLi-2 rescues Hedgehog signaling in mouse R1441C LRRK2 striatal cholinergic interneurons. (A) Confocal images of sections of dorsal striatum from 5-month-old WT or G2019S mice subjected to in situ hybridization using a PTCH1 RNA probe (white dots). Cholinergic interneurons were labeled as in Fig. 1. (B) Average number of PTCH1 dots for all ChAT^+^ cholinergic interneurons or (C) segregated according to ciliation status. Values represent mean±SEM from 4 WT and 4 G2019S brains with >45 ChAT^+^ neurons analyzed per brain. (D-F) Similar RNAscope analysis of cholinergic interneurons from R1441C LRRK2 mice fed ± MLi-2 inhibitor for 3 months. Values represent mean±SEM from 6 R1441C LRRK2 brains ± MLi-2 with >25 cholinergic interneurons scored per brain. Statistical significance was determined using an unpaired t-test or one-way ANOVA. *p =0.0195 for G2019S versus WT; ***p = 0.0001 for ciliated WT versus unciliated WT; **p = 0.0037 for ciliated G2019S versus ciliated WT. ****p < 0.0001 for + MLi-2 versus - MLi-2; ****p < 0.0001 for + MLi-2 ciliated versus + MLi-2 unciliated; ****p < 0.0001 for + MLi-2 ciliated versus - MLi-2 ciliated; **p = 0.0051 for + MLi-2 unciliated versus - MLi-2 unciliated. Bar, 10µm.

Parallel analysis revealed that striatal astrocytes from LRRK2 G2019S animals also decreased PTCH1 RNA expression upon LRRK2 mutation (Fig. 4A,B). As expected for a Hedgehog target gene, PTCH1 expression was highest in wild type ciliated cells (Fig. 4C). Upon LRRK2 mutation, even ciliated LRRK2 G2019S cells showed less than wild type PTCH1 expression.

**Fig. 4.**
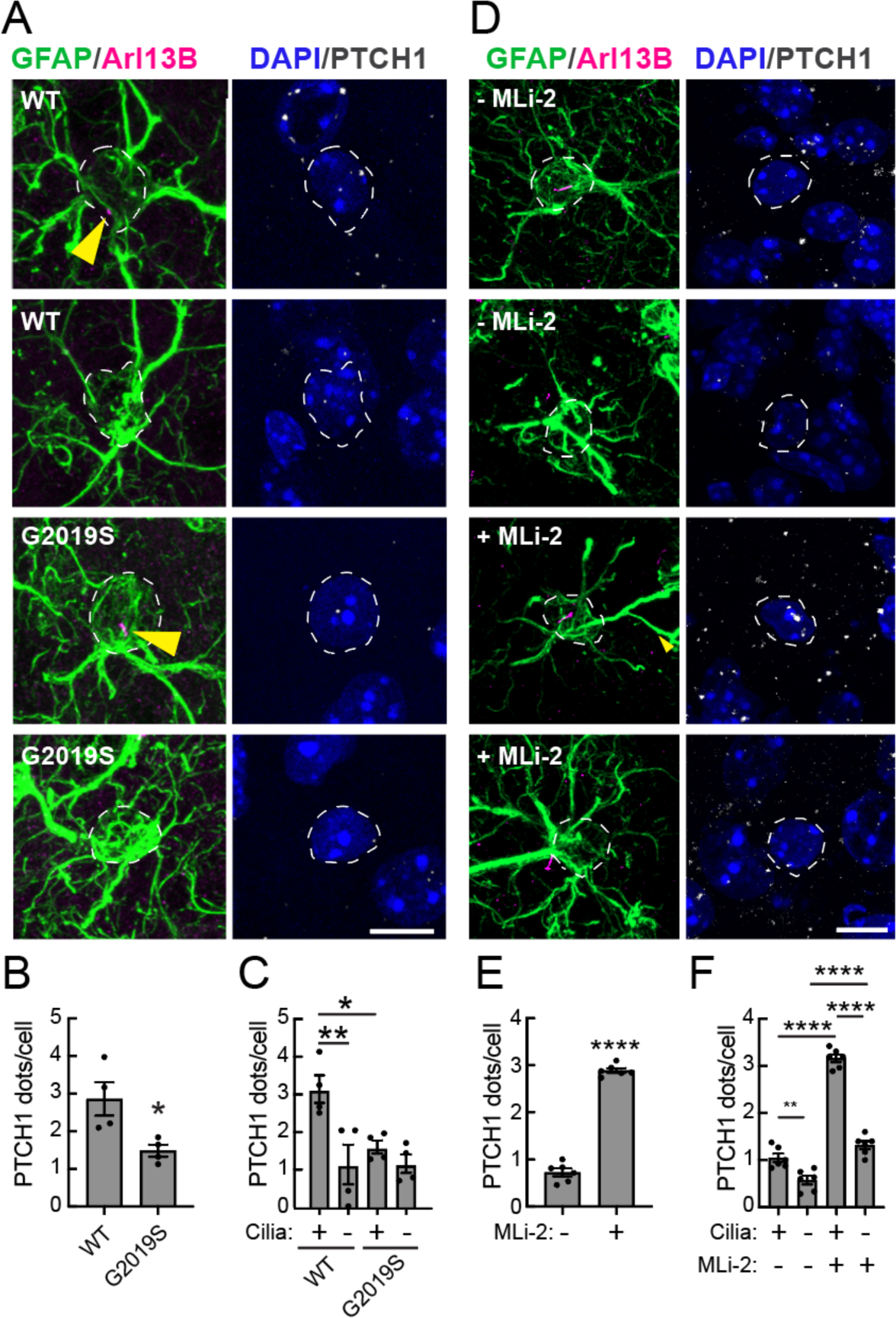
MLi-2 rescues Hedgehog signaling in mouse R1441C LRRK2 striatal astrocytes. (A) Confocal images of dorsal striatum from 5- month-old WT or G2019S mice subjected to in situ hybridization using a PTCH1 RNA probe (white dots). Astrocytes were labeled using anti-GFAP antibody (green); primary cilia were detected using anti-Arl13b antibody (magenta). Nuclei were labeled using DAPI (blue). (B) Average number PTCH1 dots for all GFAP^+^ astrocytes or (C) data segregated according to ciliation status. Values represent mean±SEM from 4 WT and 4 G2019S brains with >20 GFAP^+^ astrocytes analyzed per brain. (D-F) Similar analysis of astrocytes from R1441C LRRK2 mice (8 weeks old) fed ± MLi-2 for 3 months. Values represent mean±SEM from 6 R1441C LRRK2 brains ± MLi-2 with >30 GFAP^+^ astrocytes scored per brain. Statistical significance was determined using an unpaired t-test or one-way ANOVA. *p =0.0261 for G2019S versus WT; **p = 0.0084 for ciliated WT versus unciliated WT; *p = 0.0428 for ciliated G2019S versus ciliated WT. ****p < 0.0001 for + MLi-2 versus - MLi-2; ****p < 0.0001 for + MLi-2 ciliated versus + MLi-2 unciliated; ****p < 0.0001 for + MLi-2 ciliated versus - MLi-2 ciliated, **p = 0.0042 for – MLi-2 ciliated versus un-ciliated, ****p < 0.0001 for minus cilia, ± MLi-2. Bar, 10µm.

In similar age LRRK2 R1441C mice, MLi-2 administration increased PTCH1 expression more than 3-fold, to the level of wild type mice (Fig. 4D,E), with ciliated cells showing the greatest level of expression (Fig. 4F). Together these experiments confirm restoration of Hedgehog signaling in striatal cholinergic interneurons and astrocytes upon LRRK2 inhibition.

### Restoration of neuroprotective GDNF and NRTN production

We have shown previously that GDNF (Glial cell line-derived neurotrophic factor) and GDNF-related NRTN (Neurturin) production in cholinergic and parvalbumin neurons, respectively, is decreased in proportion to cilia loss associated with activating LRRK2 mutations (*7, 9*). This loss of protection over decades is predicted to impact the dopamine neurons that rely on these proteins for their viability. As shown in Figure 5A,B, MLi-2 administration more than doubled the production of GDNF RNA in LRRK2 mutant cholinergic interneurons, with re-ciliated cells showing the highest levels. Even non-ciliated cholinergic neurons increased basal GDNF production, with benefit for dopamine neurons (Fig 5C).

**Fig. 5.**
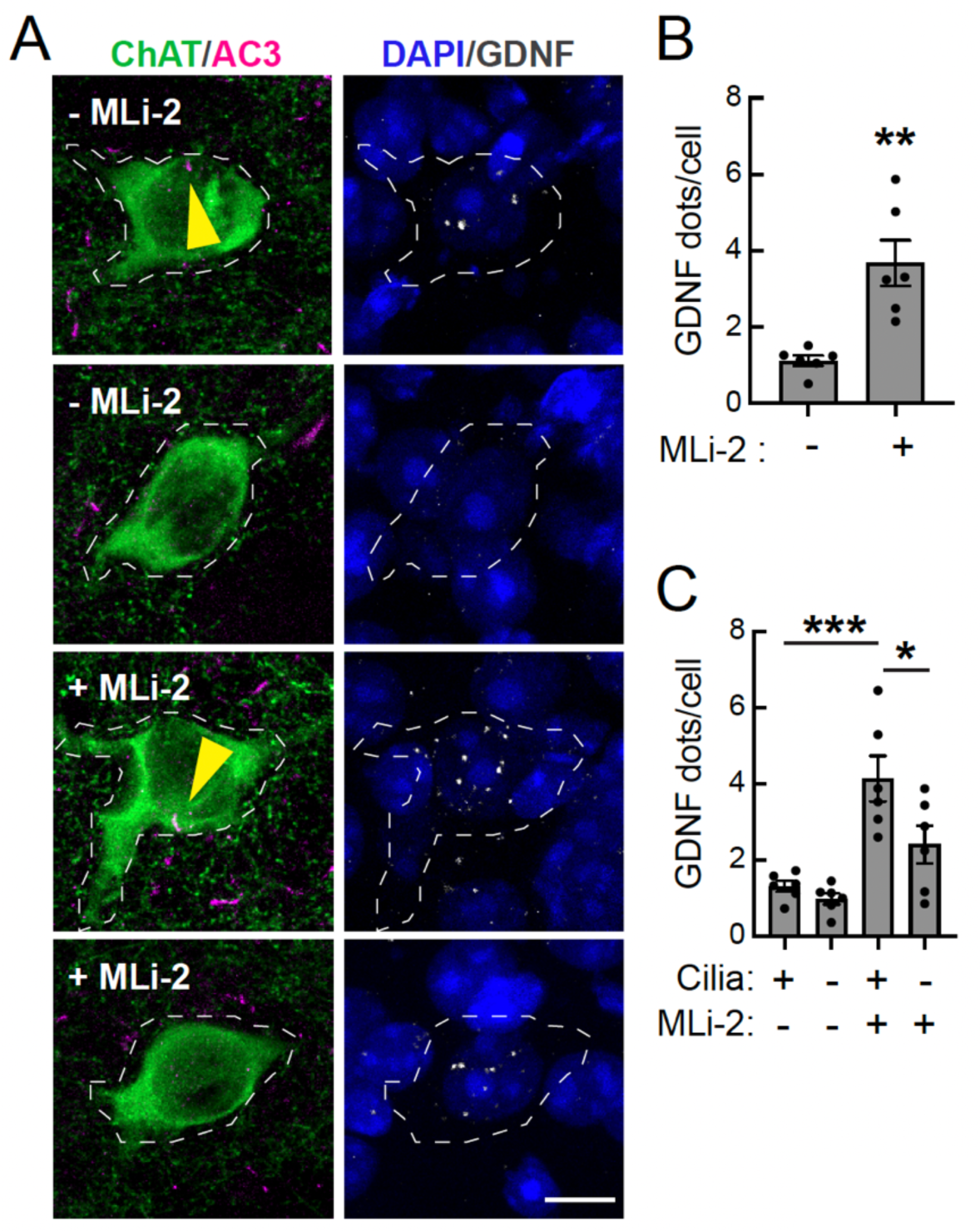
MLi-2 restores production of GDNF in mouse R1441C LRRK2 striatal cholinergic interneurons. (A) Confocal images of the sections of dorsal striatum from R1441C LRRK2 mice (8 weeks old) fed ± MLi-2 inhibitor containing chow for 3 months subjected to in situ hybridization using a GDNF RNA probe (white dots). Cholinergic interneurons were labeled as in Fig. 1. (B) Average number of GDNF dots for all ChAT^+^ cholinergic interneurons or (C) segregated according to ciliation status. Values represent mean±SEM from 6 R1441C LRRK2 brains ± MLi-2 with >25 cholinergic interneurons scored per brain. Statistical significance was determined using an unpaired t-test or one-way ANOVA. **p = 0.0019 for + MLi-2 versus - MLi-2; *p = 0.0258 for + MLi-2 ciliated versus + MLi-2 unciliated; ***p = 0.0003 for + MLi-2 ciliated versus - MLi-2 ciliated. Bar, 10µm.

Striatal parvalbumin neurons recovered their cilia to wild type levels with MLi-2, increasing from ∼55% ciliation to ∼70% ciliation, and the cilia doubled in length, consistent with reversal of cilia loss we have reported elsewhere (*9*; Fig. 6A-C). NRTN RNA was restored to a level proportional to the extent of ciliary recovery and lengthening (Fig. 6D,E), although the level in the absence of MLi-2 was higher than what we reported previously for 5-month-old LRRK2 G2019S mice; ciliated cells showed the highest level of NRTN expression, consistent with a Hedgehog-responsive process (Fig. 6E).

**Fig. 6.**
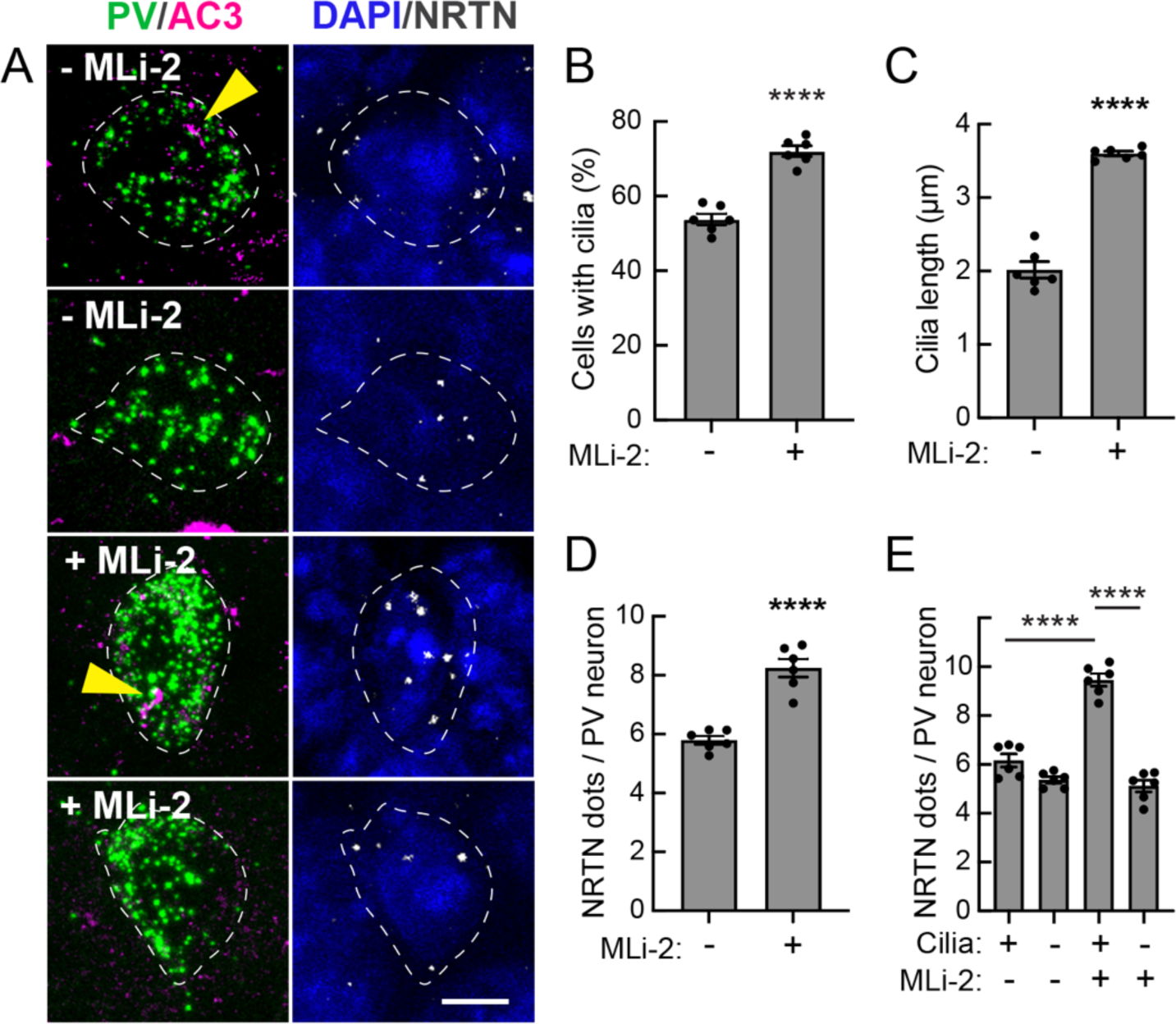
MLi-2 reverses cilia loss and restores production of Neurturin in mouse R1441C LRRK2 striatal parvalbumin interneurons. (A) Confocal images of dorsal striatum sections from R1441C LRRK2 mice fed with or without MLi-2 inhibitor chow for 3 months. RNAscope analysis of Parvalbumin (green dots) and Neurturin (NRTN) RNA (white dots) according to ciliation status (AC3, magenta). Nuclei were labeled using DAPI (blue). (B) Quantitation of the percentage of Parvalbumin neurons containing a cilium and (C) their cilia length. (D) Average number of NRTN dots for all Parvalbumin neurons or (E) segregated according to ciliation status. Values represent mean±SEM from 6 R1441C LRRK2 brains ± MLi-2 with >25 Parvalbumin interneurons scored per brain. Statistical significance was determined using an unpaired t-test or one-way ANOVA. Percentage ciliation: ****p < 0.0001 for + MLi-2 versus - MLi-2, Cilia length: ****p < 0.0001 for + MLi-2 versus - MLi-2, NRTN RNA: ****p < 0.0001 for + MLi-2 versus - MLi-2; ****p < 0.0001 for + MLi-2 ciliated versus - MLi-2 ciliated; ****p < 0.0001 for + MLi-2 ciliated versus + MLi-2 unciliated. Bar, 10µm.

### Recovery and decreased Hedgehog production by nigral dopamine neurons

In previous work we detected ∼20% pathogenic LRRK2-dependent loss of density of dopaminergic processes in the striatum, as determined by anti-tyrosine hydroxylase and anti-GDNF-receptor alpha staining of tissue sections, normalizing the staining to that detected in the same sections using antibodies that recognize total neuronal NeuN (neuronal nuclear antigen biomarker) (*7*). By this method, both tyrosine hydroxylase and GDNF receptor alpha staining were restored to wild type levels upon MLi-2 administration to LRRK2 R1441C mice (Fig. 7A-C). Strikingly, the overall density of dopaminergic processes in the dorsal striatum essentially doubled (Fig. 7D).

**Fig. 7.**
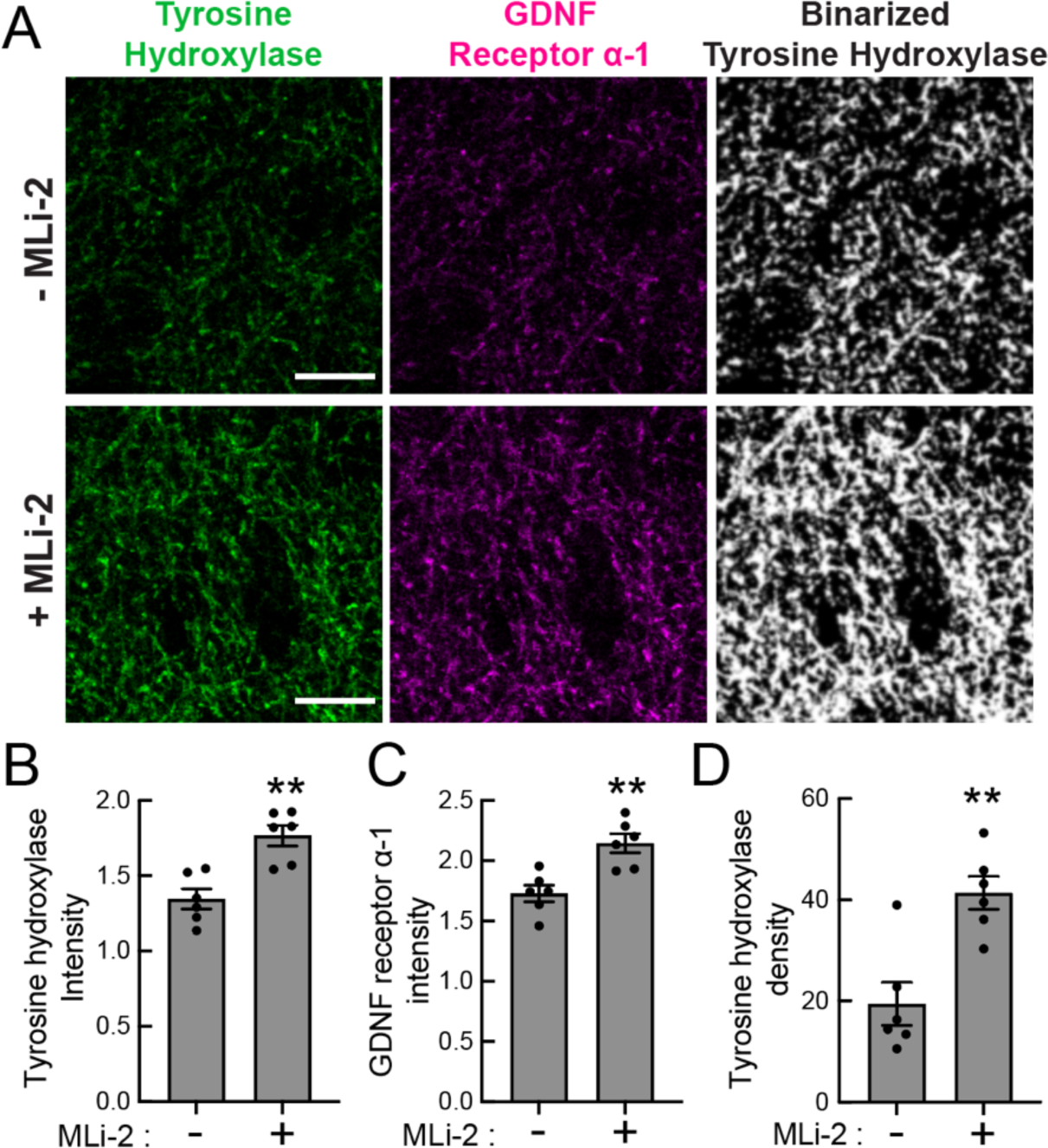
MLi-2 restores dopaminergic tyrosine hydroxylase and GDNF receptor a-1 staining in the striatum. (A) Confocal images of striatal sections from R1441C LRRK2 mice fed ± MLi-2 inhibitor, labeled using anti-tyrosine hydroxylase antibody (green) and anti-GDNF receptor alpha-1 antibody (magenta). (B) Intensity of Tyrosine hydroxylase and (C) GDNF receptor a-1 was quantified and normalized to NeuN staining using CellProfiler. Values represent mean ± SEM from 6 R1441C LRRK2 brains ± MLi-2 with >30 fields analyzed per brain. (D) Density of Tyrosine hydroxylase staining determined from binarized tyrosine hydroxylase images using a 63X objective. Each dot represents the mean of 30, 101.4 × 101.41 µm fields from a single mouse brain. Statistical significance was determined by unpaired t-test. Intensity of Tyrosine hydroxylase: **p = 0.0013 for + MLi-2 versus - MLi-2, intensity of GDNF receptor a-1: **p = 0.0026 for + MLi-2 versus - MLi-2 and *p = 0.0145 for + MLi-2 versus - MLi-2. **p = 0.0021 for + MLi-2 versus - MLi-2 density. Bar, 10µm.

It has been proposed that stress increases Hedgehog signal production by dopamine neurons (*20*). Thus, we monitored the expression of Sonic Hedgehog in dopamine neurons of the substantia nigra. In 10-month-old mouse nigra, we detected a 44% increase in Sonic hedgehog RNA upon LRRK2 R1441C mutation compared with wild type animals (Fig. 8A,B, 10-month-old mice). In our MLi-2 treatment study that employed 5-month-old mice, Sonic Hedgehog expression decreased more than two-fold, and was restored to wild type levels (at least compared with older mice) (Fig. 8C,D). These experiments confirm a profound effect of LRRK2 inhibitors throughout the nigrostriatal circuit and beyond, to the pedunculopontine nucleus.

**Fig. 8.**
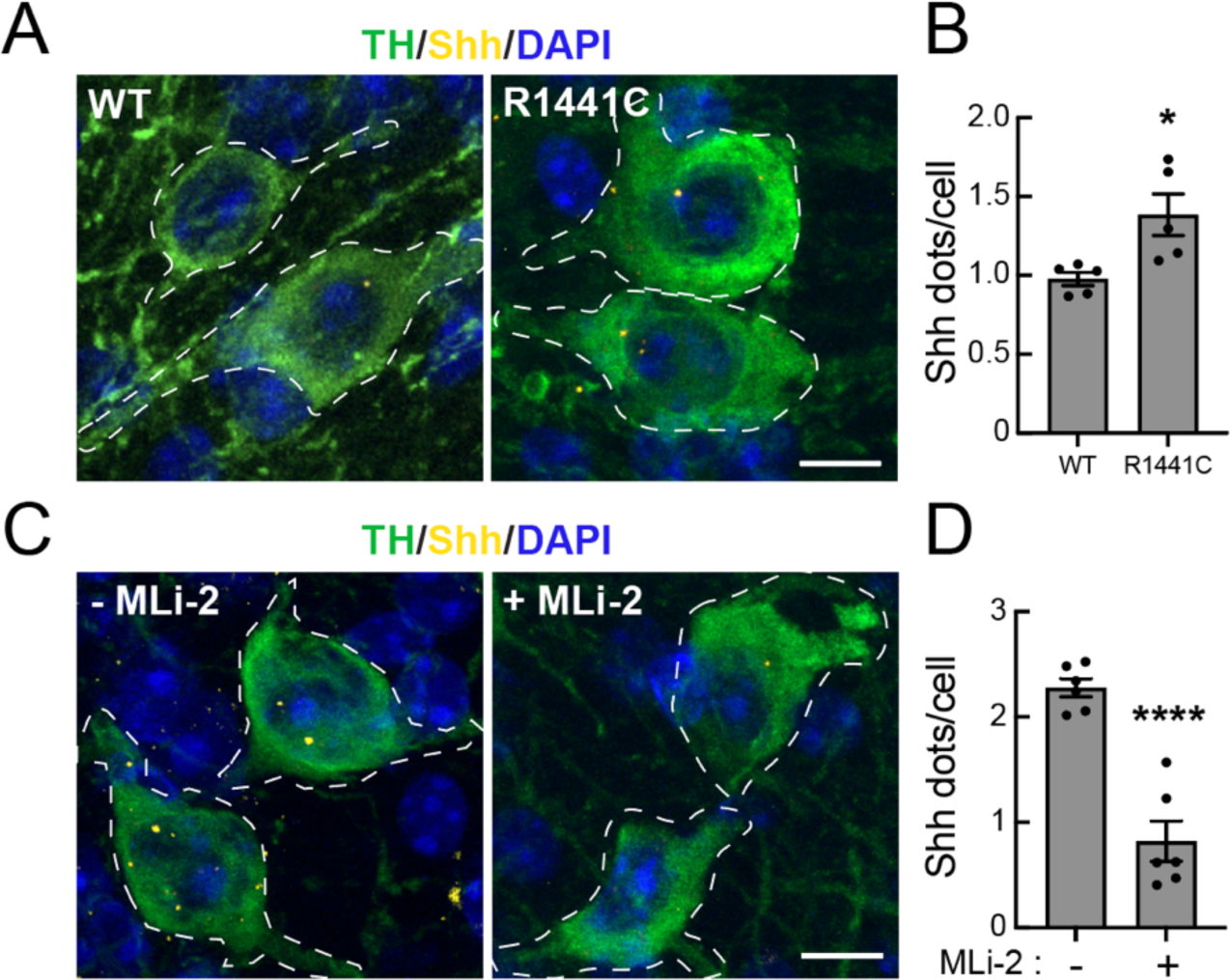
Dopamine neurons of the Substantia nigra show decreased Sonic Hedgehog expression after MLi-2 treatment. (A) Confocal microscopy of the substantia nigra pars compacta from 10- month-old WT and R1441C LRRK2 mice subjected to in situ hybridization using Shh RNA probe (yellow dots). Dopamine neurons were labeled using anti-TH antibody (green) and nuclei were labeled using DAPI (blue). (B) Quantitation of the average number of Shh RNA dots per dopamine neuron. Values represent mean±SEM from 5 R1441C LRRK2 brains ± MLi-2 with >50 dopamine neurons scored per brain. (C) Similar analysis of dopamine neurons from R1441C LRRK2 mice fed chow with or without MLi-2 inhibitor for 3 months. (D) Quantitation of the average number of Shh RNA dots per dopamine neuron. Values represent mean±SEM from 6 R1441C LRRK2 brains ± MLi-2 with >50 dopamine neurons scored per brain. Statistical significance was determined using an unpaired t-test. *p = 0.0188 for R1441C versus WT; ****p < 0.0001 for + MLi-2 versus - MLi-2. Bar, 10µm.

## DISCUSSION

We have reported here a remarkable reversal of ciliary defects in the nigrostriatal circuit of LRRK2 mutant mice. In addition to the return of cilia on striatal cholinergic and parvalbumin interneurons and astrocytes, as well as cholinergic neurons of the pedunculopontine nucleus, we measured a return of Hedgehog signaling that produced neuroprotective GDNF and NRTN proteins in the striatum. Dopaminergic processes in the striatum returned to wild type density and dopamine neurons of the substantia nigra synthesized less Shh RNA, consistent with less stress. All these changes are encouraging for patients who may be eligible for LRRK2 inhibitor therapy in the future.

It was not clear in advance that 3 months of MLi-2 feeding would reverse all ciliary phenotypes, as our previous two-week feeding regimen yielded no ciliation changes within that time window. It was possible that non-mitotic neurons and astrocytes would not be able to regrow their cilia, as cilia growth is generally a cell cycle-linked process (*12, 13*).

The possibility that at least certain post-mitotic neurons might have the capacity to regrow cilia is highlighted by a recent study of neurons that regulate our circadian clock (*21*). These authors showed that in a specific subset of suprachiasmatic nucleus neurons, cilia grow and shrink with a 12 hour period, and these rhythmic cilia changes drive oscillations of Shh signaling and clock gene expression. Thus, postmitotic cells are capable of regenerating cilia, and the cells that we study in the dorsal striatum and pedunculopontine nucleus of LRRK2 mutant mice do so on a time scale >2 weeks and < 3 months after MLi-2 administration. [Note that all the brains studied herein were harvested at a similar time of day and thus differences do not reflect possible differences in clock regulation.] Other recent metabolic studies have revealed that neuronal cilia contain both very long lived, basal body proteins as well as much shorter-lived proteins that comprise the axoneme (*22*). These disparate features suggest that cilia are indeed more dynamic than previously realized.

We showed previously that in cell culture, LRRK2-phosphorylated Rab10 protein binds to RILPL1 and in some way, phosphoRab10-RILPL1 complexes interfere with the recruitment of tau tubulin kinase 2 to release CP110 from the mother centriole (*6, 23*), a step required to initiate ciliogenesis (*24–26*). In brain, phosphoRab12 is more abundant than phosphoRab10, and phosphoRab12-RILPL1 complexes may be responsible for cilia blockade in this tissue. We hypothesize that LRRK2 inhibitors decrease the levels of phosphoRab proteins and restore tau tubulin kinase 2 recruitment to drive CP110 uncapping. How phosphoRab proteins interferes with the earliest stages of ciliogenesis will be an important question for future investigation; in addition, it will be important to elucidate why only certain cells in the dorsal striatum lose cilia, despite the presence of hyperactive LRRK2 kinase.

Limitations of this study include the fact that a relatively high dose of MLi-2 was employed, and the study was carried out in mice that likely reflect a very early stage of PD pathogenesis. Future studies can evaluate the LRRK2 kinase inhibitor concentration that is needed to achieve phenotype reversal. We recommend that this assay also be exploited to evaluate and benchmark future LRRK2 inhibitors as a key part of their preclinical evaluation.

## MATERIALS AND METHODS

### Reagents

MLi-2 LRRK2 inhibitor was synthesized by Natalia Shpiro (MRC Reagents and Services, University of Dundee) and was first described to be a selective LRRK2 inhibitor in previous work (*15*). For the MLi-2 in diet study, rodent diet containing MLi-2 at 360 mg per Kg was manufactured by Research diets, Inc.

### Research standards for animal studies

Mice were maintained under specific pathogen-free conditions at the University of Dundee (UK). All animal experiments were ethically reviewed and conducted in compliance with the Animals (Scientific Procedures) Act 1986 and guidelines established by the University of Dundee and the U.K. Home Office. Ethical approval for animal studies and breeding was obtained from the University of Dundee ethical committee, and all procedures were performed under a U.K. Home Office project license. The mice were group-housed in an environment with controlled ambient temperature (20–24°C) and humidity (45–55%), following a 12-hour light/12-hour dark cycle, with ad libitum access to food and water. LRRK2 R1441C knock-in mice backcrossed on a C57BL/6J background, were obtained from the Jackson laboratory (Stock number: 009346). LRRK2 G2019S knock-in mice backcrossed on a C57BL/6J background, were obtained from Taconic (Model 13940). Genotyping of mice was performed by PCR using genomic DNA isolated from tail clips or ear biopsies with genotyping confirmation conducted on the day of the experiment.

### In-diet MLi-2 administration

Littermate or age-matched R1441C LRRK2 homozygous knock-in mice were allowed to acclimate to the control rodent diet (Research Diets D01060501; Research Diets, Inc., New Brunswick, NJ) for 14 days before being placed on study. On day 1 of the study, one group (9 mice) received a modified rodent diet targeted to provide a concentration of 60 mg/kg per day of MLi-2 on the basis of an average food intake of 5 g/day (D19012904); the other group (9 mice) received an untreated diet and served as the control group. Bodyweight and food intake were assessed twice weekly. On day 91, mice were culled and tissues collected as described below.

### Mouse brain processing

Homozygous LRRK2-mutant (R1441C or G2019S of ages indicated) and age-matched wild type controls were fixed by transcardial perfusion using 4% paraformaldehyde (PFA) in PBS as described in dx.doi.org/10.17504/protocols.io.bnwimfce. Whole brain tissue was extracted, post-fixed in 4% PFA for 24 hr and then immersed in 30% (w/v) sucrose in PBS until the tissue settled to the bottom of the tube (∼48 hr). The brains were harvested in Dundee and sent with identities blinded until analysis was completed. Prior to cryosectioning, brains were embedded in cubed-shaped plastic blocks with OCT (BioTek, USA) and stored at −80 °C. OCT blocks were allowed to reach −20 °C for ease of sectioning. The brains were oriented to cut coronal sections on a cryotome (Leica CM3050S, Germany) at 16–25 µm thickness and positioned onto SuperFrost plus tissue slides (Thermo Fisher, USA).

### Mouse tissue processing for immunoblotting analysis

Three mice from each group (control diet and MLi-2 diet) were euthanized by cervical dislocation. Tissues including brain and kidney were collected and rinsed twice with cold PBS containing phosphatase and protease inhibitors (PhosSTOP, Merck #04906837001, and complete EDTA-free Protease Inhibitor Cocktail, Roche #11836170001) before being snap-frozen. Frozen mouse brains were placed on a stainless steel adult mouse brain slicer matrix (EMS #69090-C) kept on dry ice. Using this setup, 1.0-mm coronal sections were prepared. Landmarks in each slice were aligned with those in a reference atlas and the regions were excised using a cold scalpel or biopsy punch. For the immunoblotting shown in Fig. 1, an entire 1-mm coronal section from each sample was taken at approximately −2.3 mm rostrocaudal from bregma. The sections were then stored at −80 °C until further processing for immunoblotting analysis.

### Quantitative immunoblotting analysis

Tissue analysis by immunoblot to measure levels of Rab10, phospho-T73 Rab10, Rab12, phospho-S105 Rab12, LRRK2, and phospho-S935 LRRK2 was performed as described in https://doi.org/10.17504/protocols.io.bsgrnbv6. Briefly, snap-frozen tissues were thawed on ice in a tenfold volume excess of ice-cold lysis buffer containing 50 mM Tris–HCl pH 7.4, 1 mM EGTA, 10 mM 2-glycerophosphate, 50 mM sodium fluoride, 5 mM sodium pyrophosphate, 270 mM sucrose, supplemented with 1 μg/mL microcystin-LR, 1 mM sodium orthovanadate, cOmplete EDTA-free protease inhibitor cocktail (Roche), and 1% (v/v) Triton X-100 and homogenized using a Precellys Evolution system, employing three cycles of 20 s homogenization (6800 rpm) with 30 s intervals. Lysates were centrifuged at 15,000 × g for 30 min at 4°C, and supernatants were collected for subsequent Bradford protein assay and immunoblot analysis. The following primary antibodies were used: mouse anti-total LRRK2 (Neuromab N241A/34), rabbit anti-LRRK2 pS935 (UDD2 10(12), MRC Reagents and Services), rabbit anti-pT73 Rab10 (ab230261, Abcam), mouse anti-total Rab10 (0680–100/Rab10-605B11, Nanotools), rabbit anti-pS106 Rab12 (ab256487, Abcam), rabbit anti-total Rab12 (A26172, ABclonal). Primary antibody probes were detected using IRDye labeled secondary antibodies (IRDye 680LT Donkey anti-Mouse IgG; IRDye 800CW Donkey-anti-Rabbit IgG). Protein bands were acquired via near-infrared fluorescent detection using the Odyssey CLx imaging system and quantified using Image Studio Lite (Version 5.2.5, RRID:SCR_013715).

### Immunohistochemical staining

The mouse brain striatum was subjected to immunostaining following a previously established protocol (dx.doi.org/10.17504/protocols.io.bnwimfce). Frozen slides were thawed at room temperature for 15 minutes and then gently washed twice with PBS for 5 minutes each. Antigen retrieval was achieved by incubating the slides in 10 mM sodium citrate buffer pH 6.0, preheated to 95°C, for 15 minutes. Sections were permeabilized with 0.1% Triton X-100 in PBS at room temperature for 15 minutes, followed by blocking with 2% FBS and 1% BSA in PBS for 2 hours at room temperature. Primary antibodies were applied overnight at 4°C, and the next day, sections were exposed to secondary antibodies at room temperature for 2 hours. Secondary antibodies used were donkey highly cross-absorbed H + L antibodies conjugated to Alexa 488, Alexa 568, or Alexa 647, diluted at 1:2000. Nuclei were counterstained with 0.1 μg/ml DAPI (Sigma). Finally, stained tissues were mounted with Fluoromount G and covered with a glass coverslip. All antibody dilutions for tissue staining contained 1% DMSO to facilitate antibody penetration. Automated determination of the density and intensity of dopaminergic processes in the striatum was carried out as described: dx.doi.org/10.17504/protocols.io.x54v92km4l3e/v1.

### Fluorescence in situ hybridization (FISH)

RNAscope fluorescence in situ hybridization was carried out as described (https://bio-protocol.org/exchange/protocoldetail?id=1423&type=3) (*8*)). The RNAscope Multiplex Fluorescent Detection Kit v2 (Advanced Cell Diagnostics) was utilized following the manufacturer’s instructions, employing RNAscope 3-plex Negative Control Probe (#320871) or Mm-Ptch1-C2 (#402811-C2), Mm-Gdnf (#421951), Mm-Nrtn-C2 (#441501-C2), Mm-Pvalb-C3 (#421931-C3) and Mm-Shh-C2 (#314361-C2). The Mm-Ptch1-C2, Mm-GDNF, Mm-Pvalb-C3, and Mm-Nrtn-C2 probes were diluted 1:5, 1:20, 1:10, and 1:3, respectively in dilution buffer consisting of 6x saline-sodium citrate buffer (SSC), 0.2% lithium dodecylsulfate, and 20% Calbiochem OmniPur Formamide. Fluorescent visualization of hybridized probes was achieved using Opal 690 or Opal 570 (Akoya Biosciences). Subsequently, brain slices were subjected to blocking with 1% BSA and 2% FBS in TBS (Tris buffered saline) with 0.1% Triton X-100 for 30 minutes. They were then exposed to primary antibodies overnight at 4°C in TBS supplemented with 1% BSA and 1% DMSO. Secondary antibody treatment followed, diluted in TBS with 1% BSA and 1% DMSO containing 0.1 μg/ml DAPI (Sigma) for 2 hours at room temperature. Finally, sections were mounted with Fluoromount G and covered with glass coverslips.

### Microscope image acquisition

All images were obtained using a Zeiss LSM 900 confocal microscope (Axio Observer Z1/7) coupled with an Axiocam 705 camera and immersion objective (Plan-Apochromat 63x/1.4 Oil DIC M27). The images were acquired using ZEN 3.4 (blue edition) software, and visualizations and analyses were performed using Fiji (*27, 28*) and CellProfiler (*28*).

In addition to above-mentioned methods, all other statistical analysis was carried out using GraphPad Prism version 10.2.3 for Macintosh, GraphPad Software, Boston, Massachusetts USA, www.graphpad.com.

## Supplementary Materials

**Table 1.** Key Resources Table.

| Reagent type<br>(species) or resource | Designation | Source or<br>reference | Identifiers | Additional<br>information |
| --- | --- | --- | --- | --- |
| Genetic reagent<br>(Mus musculus) | B6.Cg-Lrrk2tm1.1Shn/J | Jackson<br>Laboratory | #009346,<br>RRID:IMSR_JAX:009346 | C57BL/6J<br>Background;<br>R1441C KI |
| Genetic reagent<br>(Mus musculus) | Constitutive KI Lrrk2tm4.1Arte | Taconic | #13940,<br>RRID:IMSR_TAC:13940 | C57BL/6J<br>Background;<br>G2019S |
| Chemical<br>Compound, drug | MLi-2 | MRC PPU<br>Reagents and<br>Services,<br>U. Dundee | CAS No. : 1627091-47-7 |  |
| Rodent diet | Control diet | Research Diets,<br>Inc. New<br>Brunswick, NJ | Research Diets D01060501 |  |
| MLi-2 modified<br>rodent diet | Diet containing MLi-2 at 360<br>mg/Kg. | Research Diets,<br>Inc. New<br>Brunswick, NJ | Research Diets D01060501 |  |
| Antibody | anti-Choline Acetyltransferase<br>(goat polyclonal) | Millipore | AB144P-1ML<br>(RRID:AB_2079751) | (1:200) |
| Antibody | anti-Adenylate cyclase III (rabbit<br>polyclonal) | EnCOR | RPCA-ACIII<br>(RRID:AB_2572219) | (1:10000) |
| Antibody | anti-GFAP (chicken polyclonal) | EnCOR | CPCA-GFAP<br>(RRID:AB_2109953) | (1:2000) |
| Antibody | anti-Arl13B (mouse monoclonal) | Neuromab | N295B/66<br>(RRID:AB_2877361) | (1:500) |
| Antibody | anti-Adenylate cyclase III<br>(chicken polyclonal) | EnCOR | CPCA-ACIII<br>(RRID:AB_2744500) | (1:5000) |
| Antibody | anti-GFR alpha-1/GDNF R<br>alpha-1 (goat polyclonal) | R&D Systems | AF560 (RRID:AB_2110307) | (1:500) |
| Antibody | anti-Tyrosine hydroxylase (sheep<br>polyclonal) | Novus<br>Biologicals | NB300-110<br>(RRID:AB_10002491) | (1:500) |
| Antibody | anti-NeuN(chicken polyclonal) | Millipore | ABN91<br>(RRID:AB_11205760) | (1:1000) |
| Antibody | H+L Donkey anti-goat<br>Alexa 488 | Life<br>Technologies | A11055<br>(RRID:AB_2534102) | (1:2000) |
| Antibody | H+L Donkey anti-Rabbit<br>Alexa 568 | Life<br>Technologies | A10042<br>(RRID:AB_2534017) | (1:2000) |
| Antibody | H+L Donkey anti-chicken<br>Alexa 488 | Jackson<br>ImmunoResear<br>ch Laboratories | #703-545-155<br>(RRID:AB_2340375) | (1:2000) |
| Antibody | H+L Donkey anti-mouse 568 | Life<br>Technologies | A10037<br>(RRID:AB_11180865) | (1:2000) |
| Antibody | H+L Donkey anti-sheep<br>Alexa 488 | Life<br>Technologies | A-11015<br>(RRID:AB_2534082) | (1:2000) |
| Antibody | H+L Donkey anti-goat<br>Alexa 568 | Life<br>Technologies | A-11057<br>(RRID:AB_2534104) | (1:2000) |
| Antibody | H+L Donkey anti-chicken Alexa<br>647 | Jackson<br>ImmunoResear<br>ch Laboratories | #703-605-155<br>(RRID:AB_2340379) | (1:2000) |
| Antibody | LRRK2 (C-terminus) | Antibodies<br>Incorporated/N<br>euroMab | 75-253<br>(RRID:AB_10675136) | (1:1000) |
| Antibody | LRRK2 pSer935 | MRC PPU<br>Reagents and<br>Services,<br>University of<br>Dundee | UDD2 10(12)<br>(RRID:AB_2921228) | (1 µg/ml) |
| Antibody | Rab10 pThr73 | Abcam Inc. | ab230261<br>(RRID:AB_2811274) | (1:1000) |
| Antibody | Rab10 | Nanotools | 0680-100/Rab10-605B11<br>(RRID:AB_2921226) | (1:500) |
| Antibody | Rab12 pSer106 | Abcam Inc. | ab256487<br>(RRID:AB_2884880) | (1:1000) |
| Antibody | Rab12 | ABclonal | A26172<br>(RRID pending) | (1:20,000) |
| Antibody | IRDye 800CW Donkey anti-Rabbit IgG | LI-COR | 926-32213<br>(RRID:AB_621848) | (1:20,000) |
| Antibody | IRDye 680LT Donkey anti-Mouse IgG | LI-COR | 926-68 022<br>(RRID:AB_10715072) | (1:20,000) |
| Commercial assay or<br>kit | RNAScopeMultiplexFluorescent<br>Reagent Kit v2 | Advanced Cell<br>Diagnostics | #323100 |  |
| Commercial assay or<br>kit | RNAScope Probe- Mm-Ptch1-C2 | Advanced Cell<br>Diagnostics | #402811-C2 | (1:5) |
| Commercial assay or<br>kit | RNAScope Probe- Mm-Gdnf | Advanced Cell<br>Diagnostics | #421951 | (1:20) |
| Commercial assay or<br>kit | RNAScope Probe- Mm-Nrtn-C2 | Advanced Cell<br>Diagnostics | #441501-C2 | (1:3) |
| Commercial assay or<br>kit | RNAScope Probe- Mm-Pvalb-C3 | Advanced Cell<br>Diagnostics | #421921-C3 | (1:10) |
| Commercial assay or<br>kit | RNAScope Probe- Mm-Shh-C2 | Advanced Cell<br>Diagnostics | #314361-C2 |  |
| Commercial assay or<br>kit | OPAL 690 REAGENT PACK | Akoya<br>Biosciences | FP1497001KT |  |
| Commercial assay or<br>kit | OPAL 570 REAGENT PACK | Akoya<br>Biosciences | FP1488001KT |  |
| Software, Algorithm | FIJI | <u>PMID:2918716</u><br><u>5</u> | <u>RRID:SCR_002285</u> |  |
| Software, Algorithm | CellProfiler version 4.2.6 | PMID:2996945<br>0 | RRID:SCR_007358 |  |
| Software, Algorithm | Graphpad Prism | Prism 10<br>version 10.2.3 | RRID:SCR_002798 |  |
| Software, Algorithm | ZEN | Zeiss ZEN<br>Microscopy<br>Software | RRID:SCR_013672<br>www.zeiss.com/microscopy/en/products/software/zeiss-zen.html |  |
| Software, Algorithm | Image Studio Lite (version 5.2.5) |  | RRID:SCR_013715). |  |

**Supplemental Fig. 1.**
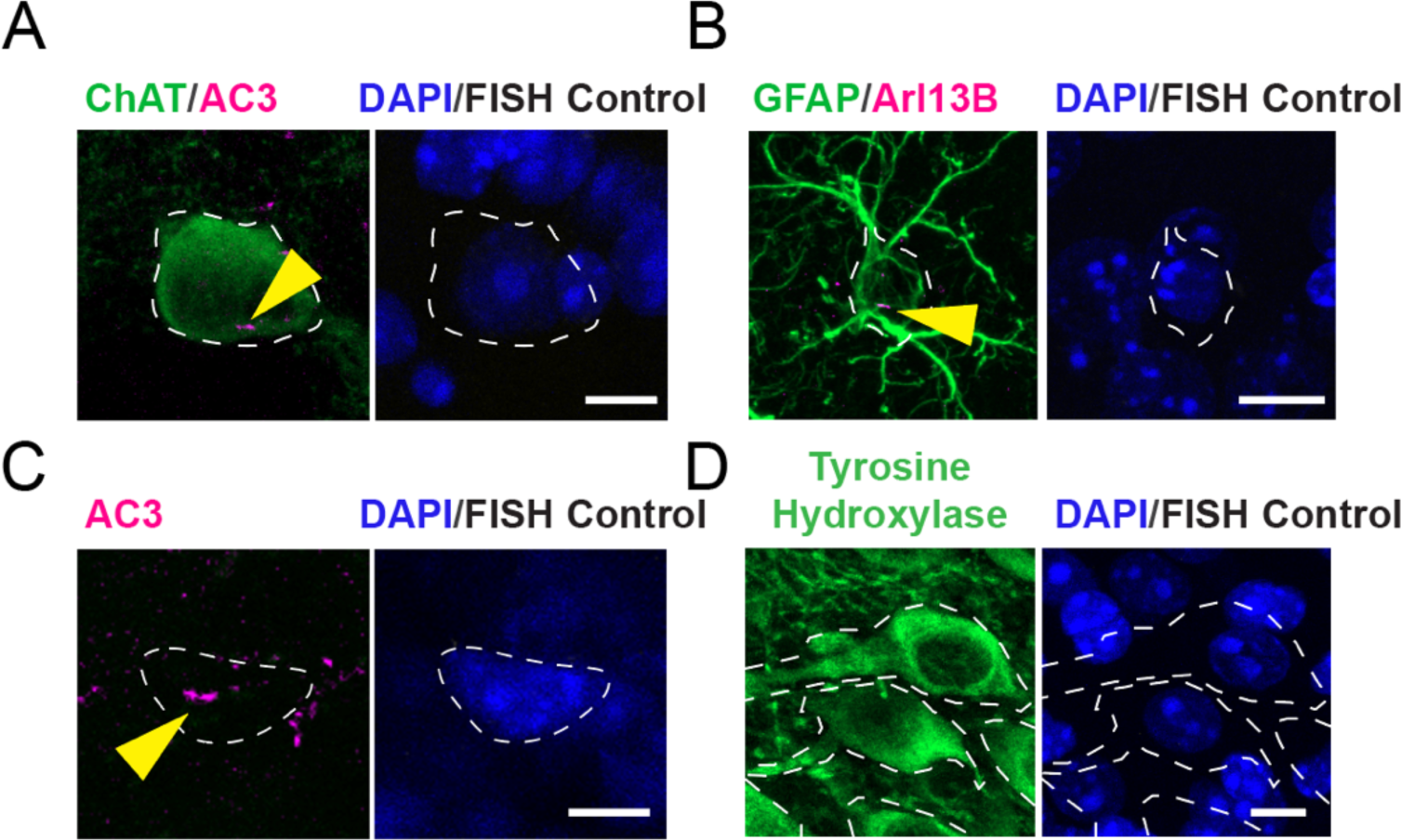
Images of control reactions carried out without the RNAScope probe for (A) cholinergic interneurons, (B) astrocytes, (C) parvalbumin interneurons, and (D) nigral dopaminergic neurons. Cholinergic interneurons, astrocytes, parvalbumin interneurons, and their cilia were labeled as described in Figs. 1 and 6. Nigral dopaminergic neurons were labeled as described in Fig 8. Bars, 10µm.

## Acknowledgments

This study was funded by the joint efforts of The Michael J. Fox Foundation for Parkinson’s Research (MJFF) and Aligning Science Across Parkinson’s (ASAP) initiative. MJFF administers the grant (ASAP-000463) on behalf of ASAP and itself (to DRA and SRP).

The DRA laboratory is also supported by the UK Medical Research Council (grant number MC_UU_00018/1).

For the purpose of open access, the authors have applied for a CC-BY public copyright license to the Author Accepted Manuscript version arising from this submission.

## Author contributions

Conceptualization: EJ, YL, DRA, SRP

Resources: FT, OA

Methodology: EJ, YL

Investigation: EJ, YL, FT

Formal Analysis: EJ, YL, FT, SRP

Visualization: EJ, YL, FT, SRP

Funding acquisition: DAR, SRP

Project administration: FT

Supervision: DAR, SRP

Writing – original draft: SRP

Writing – review & editing: SRP, DAR, EJ, YL

The authors declare that they have no competing interests.

All primary data is available at DOI: 10.5061/dryad.q2bvq83tn

